# Three-dimensional cell geometry controls excitable membrane signaling in *Dictyotelium* cells

**DOI:** 10.1101/278853

**Authors:** Marcel Hörning, Tatsuo Shibata

## Abstract

Phosphatidylinositol (3,4,5)-trisphosphate (PtdInsP3) is known to propagate as waves on the plasma membrane and is related to the membrane protrusive activities in *Dictyostelium* and mammalian cells. While there have been a few attempts to study the three-dimensional dynamics of these processes, most studies have focused on the dynamics extracted from single focal planes. However, the relation between the dynamics and three-dimensional cell shape remains elusive, due to the lack of signaling information about the unobserved part of the membrane. Here we show that PtdInsP3 wave dynamics are directly regulated by the three-dimensional geometry - size and shape - of the plasma membrane. By introducing an analysis method that extracts the three-dimensional spatiotemporal activities on the entire cell membrane, we show that PtdInsP3 waves self-regulate their dynamics within the confined membrane area. This leads to changes in speed, orientation and pattern evolution, following the underlying excitability of the signal transduction system. Our findings emphasize the role of the plasma membrane topology in reaction-diffusion driven biological systems and indicate its importance in other mammalian systems.

---

Self-organized pattern formation is ubiquitous in nature, particularly under conditions far from thermal equilibrium. The key elements behind the pattern formation are spontaneous symmetry breaking and nonlinearity. Those key elements are also present in biological cells. In particular, signal transduction systems exhibit a variety of self-organized pattern formations, such as asymmetric protein distributions and wave propagations, which play pivotal biological roles in *E. coli* (1), yeast (2), *C. elegans* (3), and chemotactic eukaryotic cells (4). During the symmetry breaking in these systems, states change from initially homogeneous to asymmetric and they are sometimes accompanied by complex interplay between system geometry and spatiotemporal signaling. How spatiotemporal signaling is related to the geometry of cells remains elusive. An asymmetric state of cell signaling dynamics is usually formed on the plasma membrane, which has characteristics of a closed and boundary-less surface in 3D. Therefore it is essential to study the relationship between pattern formation and cell geometry, to investigate the entire plasma membrane as a system.

Similar questions have been addressed in both small and large scale systems. For small reaction-diffusion type systems (5–7), chemical waves that propagate in a one-dimensional ring exhibit a modulation in speed and phase, depending on the system size(8–10). Similar behavior was also observed in large scale systems, such as spatially constrained cardiac tissue preparates(11). But such a relation has been barely investigated in closed surfaces in 3D, such as cell membranes, because it is still methodologically challenging to extract and analyze pattern dynamics on the entire living cell surface. In this study, we present a method to extract and analyze pattern dynamics on entire cell membranes, using single *Dictyostelium* cells as the model system.

A variety of spontaneous and complex pattern formation has been reported in the chemotaxis signaling pathway of *Dictyostelium* cells. The reaction dynamics of Phosphatidylinositol (3,4,5)-trisphosphate (PtdInsP3) lipids play a pivotal role in gradient sensing of chemoattractant and actin polymerization (4). In the front region along a gradient, phosphoinositide 3-kinase (PI3K) produces PtdInsP3 from Phosphatidylinositol (4,5)-trisphosphate (PtdInsP2), while phosphatase and tensin homolog (PTEN) catalyzes the reverse reaction in the rear area, leading to an accumulation of PtdInsP3 in the cell front. However, such an asymmetric distribution of PtdInsP3 and F-actin can be generated, even in the absence of a chemoattractant gradient (12–15). A variety of self-organizing patterns have been observed on the membrane, such as propagating waves and standing waves along the cell periphery in single optical sections, i.e. along a closed line(16) and on the adhesive membrane area (14, 17). These patterns have been shown to be generated by an excitable chemotactic signaling pathway(18, 19). The pattern orientation can be easily biased by external chemoattractant gradients (20, 21). Although the signaling pathways are well understood, it is still unclear how the formation of patterns on the membrane are linked to the geometry and size of the cell membrane.

Here, we approached this problem by developing an automated computational method to localize the cell membrane and extract the corresponding PtdInsP3 lipid dynamics on the entire three-dimensional plasma membrane using Delaunay triangulation. We found that differences in cell shape, i.e. size and adhesion-mediated membrane distortion, regulate the spatiotemporal PtdInsP3 dynamics. The propagation direction of PtdInsP3 domains is biased toward the longest pathway on the cell surface, and the speed of PtdInsP3 domains depends on the size of the membrane, e.g. the average domain speed increased with increasing cell size. Our findings imply a self-regulatory effect of domain dynamics that follow basic principles seen in other excitable media, such as cardiac tissue and Belousov-Zhabotinsky (BZ) reactive medium. We successfully confirmed our findings by performing additional experiments on spatially constrained *Dictyostelium* cells that were embedded in narrow grooves of micro-chambers.

## Results

### PtdInsP3 domain formation in single focal planes

Pattern formation in the PtdIns lipid signaling system of *Dictyotelium* cells was studied by measuring the distribution of PtdInsP3 using the EGFP-labeled PH-domain of Akt/PKB (PH-GFP) as shown in Fig. 1. The cells were treated with 10 *μ*M latrunculin A (an actin polymerization inhibitor) to avoid the effects of motile and protrusive activities of the actin cytoskeleton and with 4 mM caffeine to inhibit cell-to-cell interactions via secreted cAMP (16). We first observed the pattern dynamics along the cell periphery at a single focal plane, reproducing the phenomena reported previously (14, 16, 22). Typical examples are shown in Fig. 1. Fig. 1*A* shows a traveling wave that changes propagation direction spontaneously, Fig. 1*B* a more complex pattern of periodic wave generation and annihilation, and Fig. 1*C* a localized domain pattern that appears and disappears at random positions. Chaotic wave dynamics that suddenly changes to a stationary localized spot is shown in Fig. 1*D*. Though these patterns have been shown in previous studies, they give only a glimpse of the complex dynamics on the entire membrane. To get deeper insights into the formation, dynamics and influence of the membrane geometry, we have developed an analysis method to extract the pattern dynamics and cell shape in three-dimensional (3D) space.

**Fig. 1.**
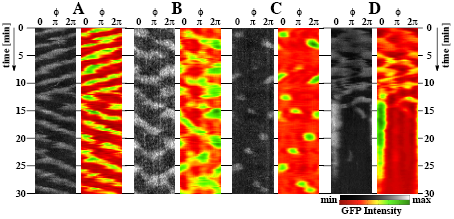
PtdInsP3 Dynamics of *Dictyostelium* cell membranes. (*A*) Propagating wave, (B) Complex wave pattern (*C*) Transient spot pattern, (*D*) Stationary spot. (*A-D*) two kymographs - raw (black-white) and analyzed (red-green) intensity data - that show the dynamics at the equatorial plane (*θ ≈*0°).

### 3D Reconstruction of Membrane Geometry and Signals

In order to extract the precise spatiotemporal pattern dynamics on the entire cell membrane, we employed a method that uses the signaling itself to specify the membrane position. Once the membrane is correctly localized, the patterns can be mapped (SI Materials and Methods).

First, the position of the entire membrane is determined. Around the cell center, grids are generated on a closed surface, such as a spherical surface, using a Delaunay triangulation routine in MATLAB (Mathworks)(23). Each grid element is identified by the azimuthal angle *ϕ*(latitude) and the polar angle *θ*(longitude) using spherical coordinates. We then considered the fluorescence intensity along the radial distance from the cell center. Along the radial distance of a grid point, the cell membrane is located at the position where the fluorescence variation is highest. This is possible because of the stable cell membrane position and the fact that the phosphorylation activity of PtdInsP2 to PtdInsP3 and vice versa is highest at the cell membrane. This procedure was independently performed for all grid elements, so that the entire membrane position could be determined and reconstructed in 3D (Fig. 2*A*).

**Fig. 2.**
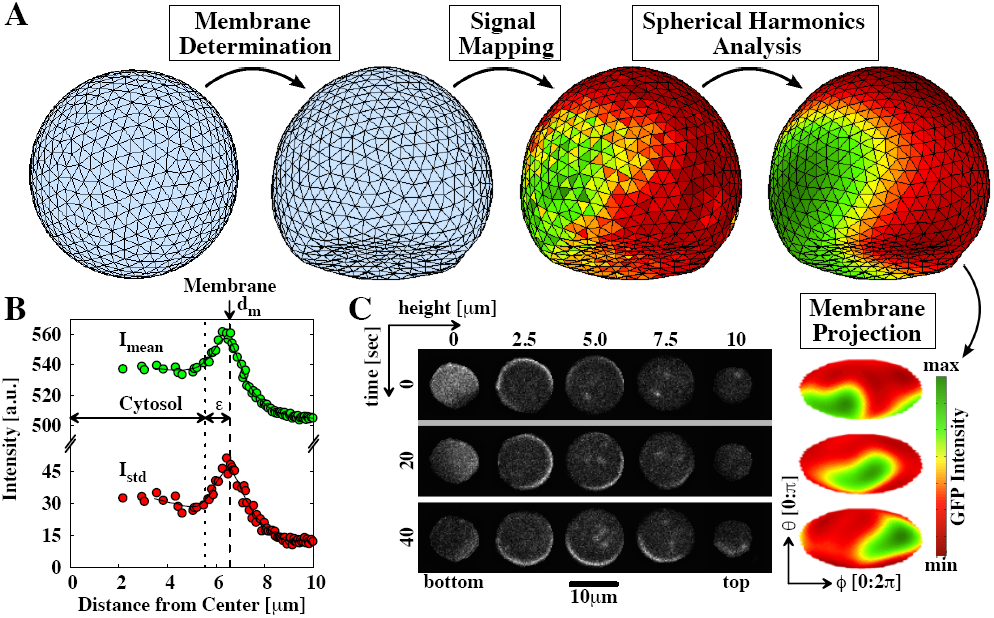
Signal analysis of PtdInsP3 waves on *Dictyostelium* membranes. (*A*) Mapping strategy of propagating waves on the entire cell membrane (Viewpoint: *ϕ* = 360 °, *θ* = *-*20 °). (*B*) The intensity profile of a single mesh unit. The upper green and lower red data points show the time-averaged intensity (*I*mean) and standard deviation (*I*_std_) of the signaling as a function of the distance from the cell center, respectively. The membrane position, *d*_*m*_ is obtained by the local maximum of *I*_std_. The mean cytosolic intensity is obtained by considering an offset ε = 1*μ*m from *dm*. (C) Raw-data snapshots at three time points and five different focal heights (bottom to top), and the corresponding Mollweide projection of the entire membrane after analysis. The second snapshot (*t* = 20 sec) corresponds to the example illustrated in (*A*) after spherical harmonics analysis. PH-GFP intensity conservation between the mean cytosolic and mean membrane intensity as a function of time is shown in Fig. S1. See also Supplemental movie 1.

Next, the fluorescence intensities on the membrane and in the cytoplasm were determined for each grid element (*ϕ, θ*) at a given time (Fig. 2*B*). The PtdInsP3 signaling distribution on the entire membrane was visualized with a 2D map using the Mollweide projection as shown in Fig. 2*C* (see alsoSupplemental movie 1). For use in later analysis, we performed a spherical harmonic transformation at each time frame and removed higher-frequency modes (Fig. 2C). Knowing the exact spatiotemporal information of the membrane geometry and signaling activity, we were able to correlate the spatiotemporal pattern dynamics and the membrane geometry.

### Pattern Dynamics on the Entire Membrane

The observation and analysis of PtdInsP3 dynamics on the entire membrane enables a more complete and precise view of the complex spatiotemporal dynamics that remain invisible in data from single focal planes. Figs. 3*A-D* show the dynamics of the four cells shown in the kymographs of Fig. 1. The Mollweide projections show snapshots of representative dynamics on the entire membrane (Figs. 3*A-D* left panels, see also Supplemental movies). In Fig. 3*A* (Supplemental movie 2), the PtdInsP3 domains propagate along the equatorial cell periphery (horizontal direction), as is expected from the kymograph in Fig. 1A. In contrast, the Mollweide projection in Fig. 3*B* reveals persistently traveling waves in the longitudinal (vertical) direction, despite the complex pattern dynamics found in the kymograph of a single section (Fig. 1B) (see also Supplemental movie 3).

**Fig. 3.**
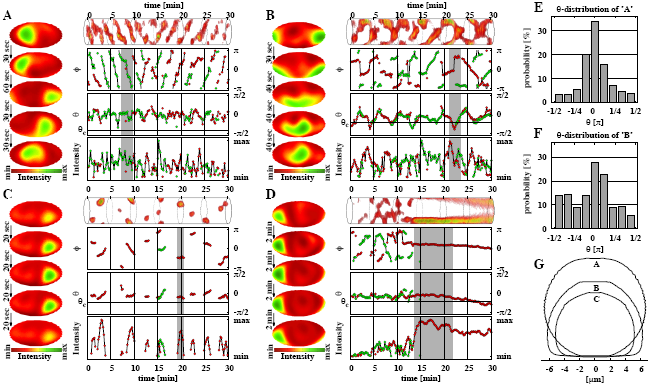
Spatiotemporal dynamics on *Dictyostelium* cell membranes. (*A-D*) Three-dimensional dynamics of the examples shown in Fig. 1, respectively. (*A*) longitudinal propagating waves (along periphery), (*B*) transversal propagating waves (bottom-top waves), (*C*) transient spot and (*D*) stationary spot. Left panel: Five prototypical subsequent snapshots. The three-dimensional kymographs show the entire membrane dynamics using the temporal integrated Mollweide projection over time (*SI Materials and Methods*). Right bottom panel: Time series of peak position given by longitude *ϕ* and latitude *θ* on the cell membrane and the corresponding intensity levels of PtdInsP3. Up to two domains are shown with red and green circles at a time. When one PtdInsP3 wave domain is present (e.g. red) the secondary wave domain is shown by the opposite color (e.g. green). The same domains are connected by a black line in time. In the time series plot of *θ*, the horizontal doted lines mark the latitude *θc* of the interface between adhesive and non-adhesive areas. Gray shaded regions correspond to the snapshots illustrated in the left panel. (*E-F*) The probability distributions of *θ* for (*A*) and (*B*), respectively. (*G*) Shape of the cells shown in (*A*), (*B*) and (*C*).

Next, we determined the precise spatiotemporal position of the domains from the local maximum PtdInsP3 intensity. We considered up to two domains at a time, because more than two domains were rarely observed and thus are statistically insignificant. The domain positions in latitude *ϕ* and longitude *θ* are plotted as a function of time (Fig. 3*A-D* right panels). Fig. 3*A* shows the domain motion in the equatorial direction, indicated by the smooth change in the latitude position *ϕ* with almost constant speed, while it was longitudinally constrained around *θ* = 0 (see Fig. 3E). Fig. 1B shows the domain motion in the longitudinal direction. A repetition of zig-zag like dynamic was found for *θ* that was accompanied by phase-jumps in *ϕ* when *θ* reached a cell pole. A comparison of the dynamics between these two cells reveals that the probability of the longitudinal direction *θ* is distributed more uniformly for the latter cell (Figs. 3E-F).

The pattern dynamics illustrated in Fig. 3C, corresponding to Fig. 1C, show domains that appear and disappear without significant propagation (see also Supplemental movie 4.). The domain lifetime was about one to two minutes. The rate of new domain formation was roughly every two to three minutes. Occasionally secondary waves formed on the opposite side of the cell membrane (see*t ∼* 15 Fig. 3*C*), indicating the existence of an inhibitory area around the first domain. This suppression of additional domains forming in the domain vicinity is commonly observed in excitable systems (24).

While most of the cells exhibited dynamic PtdInsP3 domains, only a very small fraction of cells showed stationary domains. These were high PtdInsP3 intensity, locally stable PtdInsP3 domains that lasted in some cases for more than 20 minutes, even in the absence of any external stimulus. A typical example is shown in Fig. 3D (see also Supplemental movie 5). The cell shows initially propagating waves that switch suddenly (*t∼* 13min) to a stationary domain with a much higher PtdInsP3 intensity. Although the kymograph in Fig. 1D indicates that the intensity of the domain fades out, the 3D analysis revealed that the domain drifts gradually to the bottom of the cell (Fig. 3D).

Fig. 3G illustrates the cross-sections of cell shapes along the z axis for the three cells shown in Figs. 3A-C. The first cell (A) has a large and spherical shape, while the other two cells (B and C) are distorted and thus have larger membrane areas that adhere to the substrate. In the following sections, we study how the domain dynamics correlate with the cell shape and size.

### 3D Cell Shape Distribution

We studied about 200 cells to investigate the influence of cell membrane shape on the PtdInsP3 domain dynamics. The cells were cultured on glass (*N*_glass_=79) or Polyethylenimine (PEI) coated glass (*N*_PEI_=98) to broaden the cell shape distribution (see *SI Materials and Methods*). Cell sizes vary from approximately 6 to 26 *μ*m in diameter, and 20 to 80 *μ*m in perimeter *C*_cell_. All cells were circular symmetric in the xy-section, and we characterized the cell shape by the adhesion area *A*_*adh*_ (*μ*m^2^) relative to the entire membrane area *A*_*mem*_ (*μ*m^2^), as shape parameter *r*_*a*_ = *A*_*adh*_*/A*_*mem*_ ranging from about 5 to 30 %. Without PEI the shape-distribution of *r*_*a*_ has a peak around 0 %, indicating that the cells tend to be spherical (Fig. S2D). In contrast, the peak shifts to around 15 % for cells cultured on PEI indicating more distorted shapes. This is because PEI-coated glass leads to an increase in the cell-substrate adhesion strength (25) and thus to larger *A*_adh_ with a peak at about 30*μ*m^2^ compared to 0*μ*m^2^ on glass (Fig. S2C).

### Interaction of Two Domains on 3D Cell Surface

The number of observed domains on the cell membrane is usually one and occasionally two, which also indicates that the characteristic length of the domains is comparable to the cell perimeter. The absolute distance of simultaneously occurring domains on the membrane showed a linear correlation to *C*_cell_ (Fig. 4A-B), while it did not show any significant dependence on the cell shape *r*_*a*_ (Fig. 4C). When a secondary PtdInsP3-domain appeared (Fig. 4D solid line), the angle distribution between those two domains was almost uniformly (*>* 0.3*π*). The distance between two domains increased after the appearance of the second domain, as seen in the change of the angle distribution (Fig. 4D, dashed line). We observed a shift to larger distances with a peak at around 0.65*π*. This tendency is typical in excitable systems, where the distance between two propagating waves increases until the inhibitory interaction ceases or an equilibrium is set up.

**Fig. 4.**
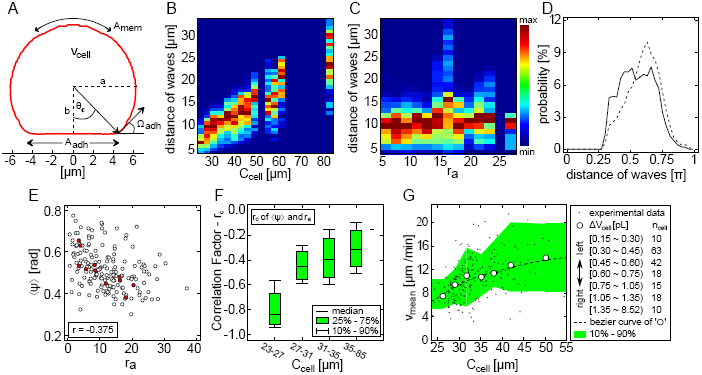
Correlation between PtdInsP3 domain dynamics and cell shape. (*A*) *ϕ*-integrated membrane profile of the cell illustrated in Fig. 2. Some of the extracted topological parameters are shown. (*BC*) Absolute peak-to-peak distance of simultaneously occurring domains on the membrane as a function of the equatorial circumferences *C*_cell_ and the shape parameter *r*_*a*_ = *A*_*adh*_*/A*_*mem*_, respectively. (*D*) The probability angle distribution of secondary PtdInsP3 domains at the time of initiation (solid line) and thereafter (dashed line). (*E*) Average direction ⟨ ψ ⟩ of domain motion as a function of *ra* for each individual cell. Red highlighted circles show cells whose circumferences are *C*_cell_ *<* 27*μ*m. (*F*) Correlation factor *r*_*c*_ calculated from (*E*) as a function *C*_cell_. (*G*) Relationship between the mean speed of PtdInsP3 domains *v*_mean_ and *C*_cell_. Each black point corresponds to a single cell. White circles are averaged data for cells of similar volume *V*_cell_, as listed in the legend box. The number of cells for each data point was set to be *n*_cell_ *≥* 10 to maintain a minimal statistical significance. The dashed line shows the Bezier curve of the white circles.

### 3D Cell Shape Constraints Dynamics of PtdInsP3 Domains

Next, we studied PtdInsP3 domain dynamics on various membrane shapes. We first considered the average direction ⟨ *ψ* ⟩ = ⟨ arctan(|*v*_*θ*_ | / |*v*_*ϕ*_ |) ⟩ of domain motions for individual cells that is defined between 0 (horizontal-direction) to *π/*2 (vertical-direction). We found that ⟨ *ψ* ⟩ is negatively correlated with the shape parameter *r*_*a*_ with a correlation coefficient of *—*0.375 (Fig. 4E). Less distorted spherical cells (*r*_*a*_ 0) are distributed around ⟨ *ψ* ⟩ ∼ *π*/4, indicating that the domain motion has no preferred direction. In contrast, as the cell shape gets more distorted (more flattened) with larger *r*_*a*_, the speed *v*_*ϕ*_ in the *ϕ*-direction tends to be larger than the speed *v*_*θ*_ in the *θ*-direction (Fig. S3C), and the domain motion tends to be more oriented to horizontal directions *ϕ*. This negative correlation is more pronounced for smaller cells (*C*_cell_ *<* 27*μ*m) (Figs. 4E-F). Thus we conclude, that the domain motion tends to orient to the horizontal direction in flattened cells.

Next, we considered the dependence of the domain speed *v*_*mean*_ on the equatorial circumference *C*_*cell*_ for individual cells. As shown in Fig. 4G, the speed tends to increase with the cell size. We sorted cells with comparable volume (*n*_cell_ *=* 10) and calculated the average domain speed *v*_mean_ for the respective cells (Fig. 4G, open circles). The obtained relationship between speed and cell size shows that *v*_mean_ increases with the increase of *C*_*cell*_. For sufficiently large *C*_*cell*_, *v*_mean_ approaches a plateau-speed of about ∼ 12*μ*m/sec, indicating that the cell size is much larger than the domain size. Such a reduction in speed has been observed in other excitable systems when the intrinsic length of a pattern is modulated (26, 27). The speed and direction of PtdInsP3 domain motion is modulated in small cells, because the intrinsic size of the domain is comparable or smaller than the length of cell surface area. In contrast, domain propagation in larger cells is less affected, i.e. the influence of the cell geometry is small. We can conclude that the membrane size and shape influences the formation, speed and propagation direction of the PtdInsP3 domains (see also Fig. S3).

### Additional Constraints in Cell Shape Modulate the Direction of Domain Motion

To strengthen the conclusion of our findings, we further modified the cell shape using a microfluidic chamber keeping the 3D analysis of the domain dynamics. We cultured *Dictyostelium* cells in 5 *μ*m wide and 10 *μ*m deep grooves, and consistently treated the cells with 20*μ*M latrunculin A and 4*m*M caffeine, as before (*SI Materials and Methods*). We considered cells with small volumes only (*V <* 0.3*p*L, *n* = 6). We expected that the shape and limited area of the membranes strongly affect the PtdInsP3 domain dynamics. Fig. 5A shows an example of such a cell. The cell forms an ellipsoidal shape that is flattened horizontally in contrast to cells that adhere to glass, i.e., *x*-axis is shortest, while circular symmetric in *yz*-plane. In Fig. 5B, we show the average direction ⟨ *ψ* ⟩ of domain motion, as introduced in Fig. 4E. As expected, we observed with this topological constraint that the PtdInsP3 domains propagate more dominantly along the longitudinal direction (Figs. 5B-C) than in cells that adhere to glass (with and without PEI). This result gives another strong indication that the 3D cell shape affects the pattern formation of PtdInsP3 in *Dictyostelium* cells.

**Fig. 5.**
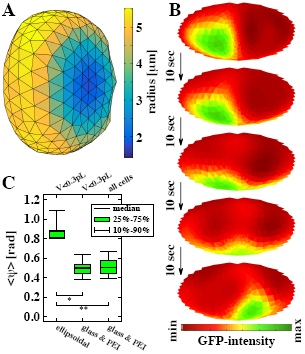
PtdInsP3 Dynamics of spatially confined *Dictyostelium* cells. (*A*) Three-dimensional map of a cell cultured in a narrow PDMS chamber. The color code depicts the radius of each grid element from the center of the cell. (*B*) Projections of typical longitudinal directed dynamics of a PtdInsP3 domain. (*C*) The median of the average direction ⟨ | ψ | ⟩of domain motion for small ellipsoidal cells (*V <* 0.3*p*L), small glass and PEI-adhered cells (*V <. p*L) and all adhered cells, respectively. The p-values were determined by two-sided Wilcoxon rank sum test, p*<*0.0005 (*) and p*<*0.00005 (**).

## Discussions

Signaling dynamics of amoeboid cells have been studied in depth, revealing intracellular mechanisms of cell migration dynamics and protrusion formation of the membrane edge in *Dictyostelium* and other mammalian cells (4). In previous studies, only a fraction of the cell membrane was typically observed, which is insufficient to understand the complex multi-dimensional signaling dynamics. In the present study, we developed a membrane extraction and signal mapping methodology that enables the correlation between PtdInsP3 signaling dynamics and cell morphology of the entire three-dimensional plasma membrane. We found that cell size and shape affect the spatiotemporal pattern dynamics, and that the speed of domain motion increases with increasing cell size, while in flattened cells, the direction of motion tends to be perpendicular to the flattened direction.

We performed stochastic simulations to see whether these cell topological constraint dynamics are reproducible in a reaction-diffusion type system (Fig. S4, supplemental material). We used a previously proposed model of PtdInsP3 pattern formation (16, 22) and extended it to realistically shaped three-dimensional cell surfaces. With that, we were able to reproduce the main features, as we observed experimentally. In particular, the speed of domain motion was increased as the cell size increased (Fig. S4A), while the direction of motion tended to be oriented along the horizontal direction (*ϕ*-direction) (Fig. S4B-C). We also observed that the bias in the propagation direction increased with an increase in the cell shape distortion. Although the simulations capture the experimental findings well, the difference between theoretical and experimental results (Fig. S4C) imply additional, still unknown effects in real *Dictyostelium* cells.

From our results, we speculate that the PtdInsP3 domains tend to propagate in the direction of the longest possible propagation path. In the case of strongly distorted cells (large *r*_*a*_), this path is the equatorial circumference. In contrast, cells that are horizontally fixed in PDMS grooves have their longest propagation path along the vertical axis, and thus, dominantly show vertically directed wave dynamics. Because the membrane curvature along the equatorial circumference in flat cells is relatively high, and the confined cells shown in Fig. 5 show high curvature area along the vertical direction, the PtdInsP3 domain pattern may have a tendency to move along this high curvature region. Considering that the leading edges of migrating *Dictyostelium* cells is a high curvature area of flatter shape, such a tendency may be appropriate. Additional studies would be necessary to understand how different shapes of closed surfaces affect pattern dynamics.

Our observations reveal that PtdInsP3 domains can propagate in any direction depending on the shape of the 2D cell membrane. Single focal plane observation at the adhesion area has shown that pattern dynamics may appear as translational waves or self-sustaining rotational waves(14). In contrast, our results show that those two types of waves can be classified as the same type of wave with different propagation directions. Waves that propagate along the horizontal direction (*ϕ*-direction) appear as self-sustaining rotational waves at the adhesion area (Fig. 1A and Fig. 3A), whereas translational waves at the adhesion area are self-sustaining rotational waves that propagate along the vertical direction (Fig. 1B and Fig. 3B).

Most cells exhibit waves with interchanging propagation directions during the course of observation. Changes in direction from clockwise to anti-clockwise, or vice versa, are observed in the case of cells that show horizontal wave propagation dominantly (Fig. 1A, ∼12 min). During this directional change, the domain often propagates vertically for a brief moment, as indicated by the zig-zag pattern of the *θ* position (Fig. 3A, ∼12 min). Whether this change is noise-induced or mediated by other intrinsic dynamics remains to be elucidated in future investigations. However, we expect it is a general concept, because such dynamics are also seen in stochastic simulations (Fig. S4E). Additionally, we remark that the wave domains continuously propagate at the interface between adhesion and non-adhesion membrane areas without the disappearance of domains or other noticeable effects (The interface line at *θ* = *θ*_*c*_ is indicated in Fig. 3A-D and Supplmental Movies). This implies that the interface plays no role in the pattern dynamics, contrary to a previous report(14).

Many cellular subsystems of *Dictyostelium* cells, such as the actin cytoskeleton (28) and the signaling pathway(18, 19), have been reported to exhibit excitable properties. The relationship between domain speed and cell size shown in Fig. 4FG can be interpreted in terms of the dispersion relation (velocity restitution) of excitable systems. Classically, the dispersion relation indicates that the speed of waves decreases when the distance between two subsequently activated waves decreases, as observed in spatially constrained BZ media, such as spherical surfaces(5) and rings(8–10), and cardiac tissue (29). Here, we consider the equatorial circumference instead of the wave distance due to the complexity of the 2D closed membrane. Our findings suggest that the pattern dynamics adjust dynamically to changes in the surface size and shape of the membrane. In the case of small cells, the domains may fail to propagate, as shown in (Fig. 3C)(30). In the case of motile cells, we expect a decrease in the propagation speed in sufficiently small membrane regions, such as in pseudopods, leading to a domain localization that may act as an activation area of other signaling pathways. How the heterogeneity in shape and size of biological systems controls the spatiotemporal pattern formation dynamics is an intriguing future topic for in-depth studies in biophysics.

The proposed three-dimensional analysis method enables the realistic comparison of PtdInsP3 domain dynamics and membrane geometry. In this study, we take advantage of the simple geometry of actin polymerization inhibited *Dictyostelium* cells. A next challenge would be to apply this method to motile *Dictyostelium* cells. Additionally, other more complex cellular systems can be considered, such as cancer cells and rapidly beating heart cells by adding an analysis of the actin remodeling dynamics (31, 32).

## Materials and Methods

*Dictyostelium discoideum* cells were cultured according to standard procedures. Fluorescence images were obtained using an inverted microscope (IX81-ZDC2; Olympus) equipped with a motorized piezo stage and a spinning disc confocal unit (CSU-X1-A1; Yokogawa). Full methods and any associated references for cell preparations and analysis are described in SI Materials and Methods.

## ACKNOWLEDGMENTS

All authors thank M.-L. Latteyer, M.-L. Lemloh, M. Tanaka, I. Weiss and T. Bullmann for their fruitful discussions and helpful comments on the manuscript. We would also like to thank H. Miyoshi (Tokyo Metropolitan Univ.), M. Nishimura and M. Sugawara (Chiba Univ.) for providing the microfluidic chambers, and Y. Wada for the experimental support. M.H. thanks JSPS for the financial support (KAKENHI, Grant No. 16K18525).

## Supporting Information

### Supporting Information (SI)

#### Cell preparation

*Dictyostelium discoideum* cells were used to observe spatiotemporal dynamics of PH_*Akt/PKB-EGFP*_. GFP-fused fused pleckstrin-homology domain of Akt/PKB (PHAkt/PKB) was expressed in wild-type AX-2 cells. Cells were cultured at 21°C in HL-5 medium and selected with 20*μ*g*/*mL G418 (1). Before observation the cells were placed in glass bottom dishes (IWAKI, Japan) 27 mm in diameter and starved in development buffer (DB: 5 mM Na phosphate buffer, 2 mM MgSO4, and 0.2 mM CaCl2, pH 6.3) for 4.5 hours, leading to chemotactic competency with respect to cAMP(2). The glass bottom dishes were used in the presence or the absence of Polyethylenimine (PEI) at a concentration of 2.4*μ*g/mL in DB. The number of cells was maintained at less than 5 ± 10^5^ cells to sufficiently separate neighboring cells. After starvation the extracelluar fluid was exchanged and additionally 20 min incubated before observation with 1 *m*L DB supplemented with 20*μ*M latrunculin A (L5163-100UG, Sigma), which inhibits actin polymerization and adenylylcyclase, and 4*m*M caffeine, which is responsible for intrinsic cAMP production. The latter is required to observe self-organizing patterns(2). All in all 79cells were observed on glass and 98cells on PEI-coated glass. A microfluidic chamber with 5 *μ*m wide and 10 *μ*m deep grooves made of polydimethylsiloxane (PDMS, Sylgard 184 Silicone Elastomer, Dow Corning) were used. Only cells that were completely fixed between the grooves where observed and analyzed (*V* < 0.3*p*L, *n* = 6).

#### Cell observation

Fluorescence images were obtained by using an inverted microscope (IX81-ZDC2; Olympus) equipped with a motorized piezo stage and a spinning disc confocal unit (CSU-X1-A1; Yokogawa) through a 60x oil immersion objective lens (N.A. 1.35; UPLSAPO 60XO). PtdInsP3-GFP was excited by 488-nm laser diode (50mW). The images were passed through an emission filter (YOKO 520/35, Yokogawa) and captured simultaneously by a water-cooled EMCCD camera (Evolve, Photometrics). Time-lapse movies were acquired at 10-s intervals at a spatial resolution of dx = dy = 0.2666*μ*m and dz = 0.5*μ*m using z-streaming (MetaMorph 7.7.5). Cells observed in the microfluidic chamber are acquired at a spatial resolution of dx = dy = 0.2222*μ*m and dz = 0.2*μ*m.

#### 3D Reconstruction of Membrane Topology and Signals

In order to map and analyze the entire *Dictyostelium* membrane topology and PtdInsP3 signaling dynamics, we considered the spatial information as a matrix consistent of all three spatial directions (*x, y* and *z*), rather than analyzing recorded focal heights (*x* and *y*) independently. A spherical grid was generated and centered to that of the cell using a Delaunay triangulation routine in MATLAB (Mathworks) (3) (Fig. 2A). The spatial coordinates were sorted to the tetrahedral shaped grid elements, as **M**_m_(*x, y, z*), considering the corners of each tetrahedron to be the center of cell and the triangular membrane node. The intensity profile for each **M**_m_ is calculated as the time-averaged standard deviation as a function of the distance from the center, *d*. The membrane position of a grid is determined at the position *d* = *d*_*m*_ with the maximum Intensity at *d*_*m*_. The number of nodes, *m*_*max*_, is then iteratively determined as a function of the respective cell size, observe spatiotemporal dynamics of PH_*Akt/P* *KB*_-EGFP. GFP-so that the average area, *A*_*m*_, of all nodes at a given mesh was approximately 0.3*μm*^2^. The number of grids ranged from *m*_*max*_ = 620 to 6632 nodes that correspond to the smallest and largest observed cell, respectively. Figure 2B shows an example of the time-averaged intensity profile *I*_mean_ (green) and the temporal standard deviation *I*_std_ (red) extracted from one tetrahedron (node). The dashed line shows the position of the membrane defined by the maximum in *I*_std_, that is set by definition as *d*_*m*_. We have chosen to use *I*_std_ instead of *I*_mean_ to define the membrane position at each **M**_m_, because the fluorescence signals on the membrane and in cytosol can overlap leading to an underestimation of *d*_*m*_. Additionally, *I*_std_ is the more sensitive marker in case of cells that exhibit low PtdInsP3 activity. Thereafter, *d*_*m*_ is smoothed among neighboring nodes using a triangular low-gaussian kernel as

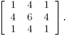

Finally, the membrane and cytosolic intensities for each node is extracted at a given time. An offset of *Á* = 1*μm* is considered to separate the cytosolic intensities from the membrane intensity (see Fig. 2B). The effect of bleaching is corrected considering a single exponential decay that fitted the data best. Fig. S1 shows the GFP-intensity conservation between the mean cytosolic and mean membrane intensity as a function of time for the cell illustrated in Fig. 2*A*.

We used the spherical harmonic expansion to smoothen the intensity distribution for peek detection. The membrane intensity *I*_mem_ (*ϕ, θ*) is expanded in the spherical harmonics as

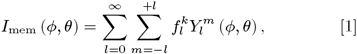

where the spherical harmonics is given by

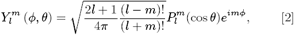

with the Legendre Polynoms 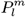. The membrane intensity is reconstructed by removing higher frequency modes as,

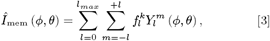

In this study *l*_*max*_ = 3 is used that corresponds to the octupole moment. To determine the location of PtdInsP3 intensity domains, 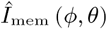 was used.

#### Mollweide Projection

The Mollweide projection is a homolo-graphic equal-area projection (4), i.e. the area accuracy based projection, which is generally used for visualizing global distributions (5), such as the cosmic microwave background and cerebral cortex(6). The transformation from spherical co-ordinates (*ϕ, θ*) to cartesian coordinates (*x, y*) is performed through the equations

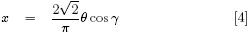

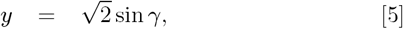

with the auxiliary angle *γ* is given as

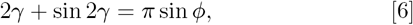

which can be solved iteratively by the Newton-Raphson method (4, 7). For simplicity only the nodes of the three-dimensional triangular grid have been projected leading to a non-smoothened periphery of the projection area (i.e. Fig. 2C).

#### Estimation of adhesion energy

The adhesion energy *E* of cells on PEI coated glass, *E*_PEI_ = 0.75 ± 0.40*n*N*μ*m^−1^, was larger than that for cells on glass, *E*_glass_ = 0.37 ± 0.30*n*N*μ*m^−1^. *E* is calculated by the Young-Dupre Equation (8) assuming tension-dominant adhesion of cells (plastic deformation only) (9, 10). *E* is calculated as follows

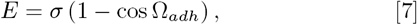

where σ = 1*n*N*μ*m^−1^ is the cortical tension(11), as measured in wild type *Dictyostelium* cells (12, 13).

#### Wave Peak and Speed Analysis

The wave domains (*P*_*n*_(*ϕ, θ, t*) ≤ 2) were tracked and analyzes considering a time-independent threshold *I*_thresh_ = (*I*_max_ – *I*_min_)*/*2. Wave speed on the membrane is calculated considering the displacement on the membrane between temporal subsequent peak positions, as

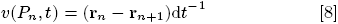

with **r**(*P*_*n*_) = (*R, θ, ϕ*), where *n* is the temporal index of the corresponding domains *P*_*n*_. The distance of simultaneously occurring waves is approximated as the shortest distance along the great-circle on the membrane

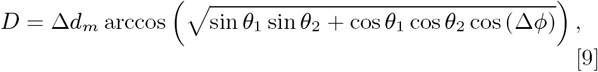

with Δ *d*_*m*_ = |*d*_*m*_,_2_ – *d*_*m*_,_1_|, Δ*θ* = |*θ*_2_ – *θ*_1_| and Δ*ϕ* = |*ϕ*_2_ – *ϕ*_1_| where the indexes correspond to the first and second peak at a given time step, respectively.

#### Numerical simulation of mathematical model

The reaction scheme of the mathematical model is given by

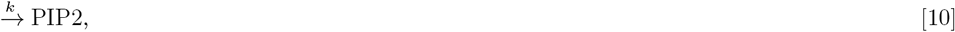

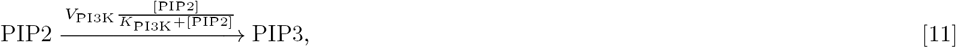

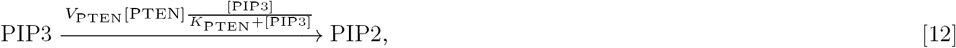

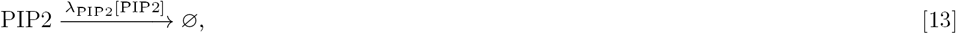

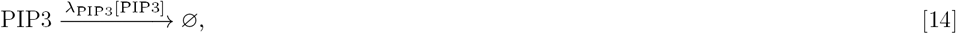

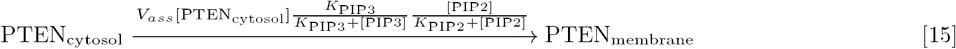

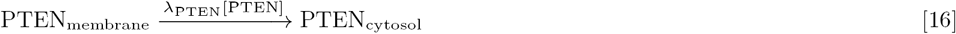

with

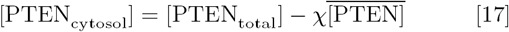

where [PIP2], [PIP3], and [PTEN] are the membrane concentrations of PtdIns(3,4,5)P3, PtdIns(4,5)P2, and PTEN, respectively, [PTEN_cytosol_] and [PTEN_total_] are the cytosolic and total concentrations of PTEN, respectively, and 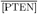 is the average membrane PTEN concentration. Further, *V*_[*protein*]_, *V*_*ass*_, *K*_[*protein*]_, and */*_[*protein*]_, *χ* are constant parameters. The diffusion of [PIP2] and [PIP3] are defined by the diffusion constants *D*_[*protein*]_. Further information of the dynamics are described in detail elsewhere (2, 14). We performed stochastic simulations on the surface of a three-dimensional cell shape using *τ*-leap method (15, 16). On the surface, triangular uniform mesh is generated (see Fig. 1A), similarly as used for the experimentally observed cells. The time stepping of the *τ*-leap method is chosen, as *τ* = 2 10^−4^ sec. The simulations are performed for a duration of 5 10^3^ sec. We varied the cell size between 4 *μ*m and 16 *μ*m by considering three different membrane shape parameter *r*_*a*_ as 0%, 20% and 30% corresponding to grids with 3608, 3548 and 3520 nodes, respectively. This captured the range of cell sizes and shapes that we observed experimentally. For each of these 39 morphologically different conditions we performed 30 simulations leading to more than 1000 independent scenarios. The obtained spatiotemporal patterns were analyzed using the same tools as for the experimental data to extract the positioning and dynamics of the PtdInsP3 domains. Figure S4 shows the results of the numerical simulations. The results obtained through the numerical simulation are analyzed using the same algorithms as the experimental observed data.

## SI Tables

**Table S1.** Parameter of Cell Morphology and Dynamics

| | wave type | $V_{\text{cell}}$<br>[pL] | $A_{\text{adh}}/A_{\text{mem}}$<br>[%] | $C_{\text{cell}}$<br>[$\mu\text{m}$] | $\Omega_{\text{adh}}$<br>[$^{\circ}$] | $v_{\text{mean}}$<br>[$\mu\text{m}/\text{min}$] | $v_{\phi}/v_{\theta}$ |
| --- | --- | --- | --- | --- | --- | --- | --- |
| A | longitudinal | 1.25 | 6.0 | 42.5 | 30 | 19.3 | 1.7 |
| B | transversal | 0.89 | 22.2 | 39.7 | 70 | 10.7 | 2.0 |
| C | spot | 0.53 | 18.6 | 32.8 | 58 | 4.84 | 1.7 |
| D | standing | 1.30 | 14.6 | 44.1 | 47 | 3.9 | 2.1 |

## SI Figures

**Fig. S1.**
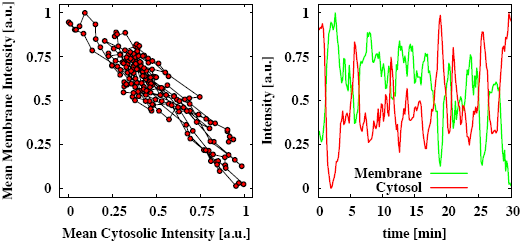
Conservation of PH-GFP intensity between the mean cytosolic and mean membrane intensity as a function of time. This dynamics correspond to the cell illustrate in Fig. 2.

**Fig. S2.**
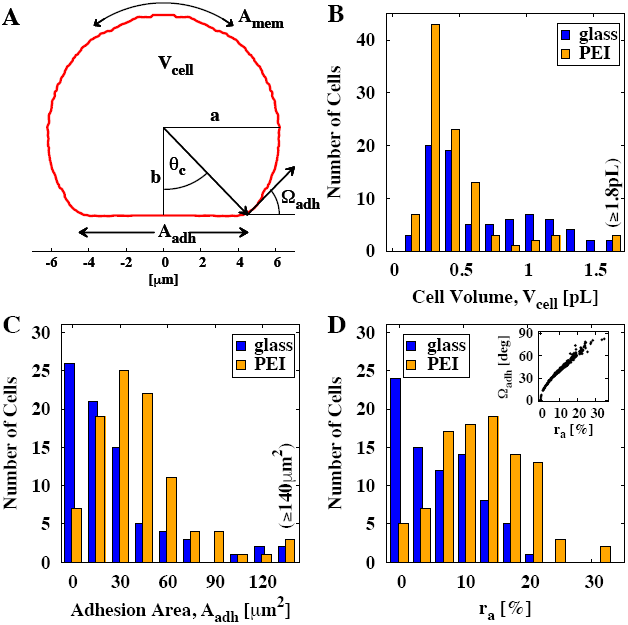
Quantification of *Dictyostelium* membrane topology. *(A)* shows exemplarily the profile (*ϕ*-integrated) of the *Dictyostelium* cell-membrane that is illustrated in Fig. 2 and some of the topological parameter that are extracted from the data. *(B-D)* show the histograms of cell volume, adhesion area and the shape parameter *r*_*a*_, respectively. The inlet in *(C)* shows the correlation between *r*_*a*_ and the contact angle n_*adh*_ of the cell. Blue and orange color depict the cells cultured on glass and PEI coated glass substrates, respectively.

**Fig. S3.**
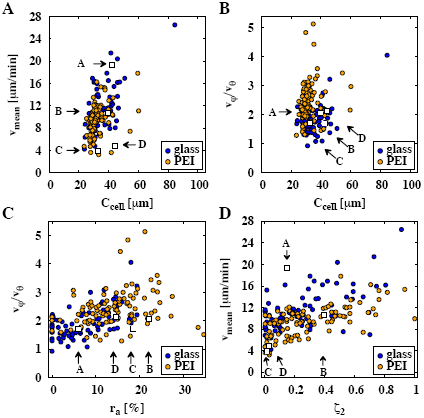
Quantification of PtdInsP3 domain signaling dynamics. *(A-D)* show wave speed and wave speed ratio (longitudinal to lateral speed) as functions of equatorial circumferences C_cell_, the shape parameter *r*_*a*_ and the probability of secondary waves, *’*2. The latter describes the probability a second wave is present during the time of record, i.e. *’*2 =0 if only single waves are detected and *’*2 =1 if second waves are always detected. Blue and orange data points depict *Dictyostelium* cells cultured on glass and PEI treated glass, respectively. The examples discussed in Figs. 1 and 3 are highlighted by white squares and respective labels from A to D.

**Fig. S4.**
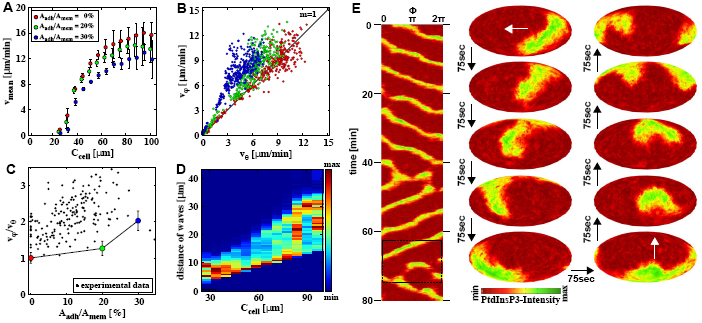
Quantification of stochastically simulated PtdInsP3 signaling dynamics. *(A)* shows average wave speed versus the cell circumferences (C_cell_). *(B)* shows average longitudinal versus lateral wave speed for each simulated cell. *(C)* shows the average wave speed ratio (longitudinal to lateral speed) as functions of aspect ratio of adhesion area and total membrane area (*A*_*adh*_*/A*_*mem*_). The ratios are obtained considering the initial slope in *(B)* only, to depict a better comparison to the experimental data (black data points, as shown in Fig. 5G. The color coding depicted in *(A-C)* corresponds to the aspect ratios of adhesion area and total membrane area (*A*_*adh*_*/A*_*mem*_) as 0% (blue), 20% (green) and 30% (red). *(D)* shows the relative occurrence of the distance of waves as a function of C_cell_. Error bars indicate the standard deviation of the data points. *(E)* shows an example simulation of PtdInsP3 signaling dynamics that exibits dominant horizontal waves (*A*_*adh*_*/A*_*mem*_ = 30%, *R*_*cell*_ = 6*μ*m). The kymograph shows the dynamics at the equatorial plane (*θ* 0*°*). The snapshots illustrate a transition from longitudinal (left column) to lateral waves (right column). The white arrows indicate the direction of motion.

## SI Movies

**Movie S1.**
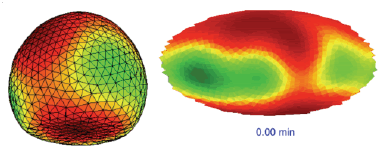
Animation of PtdInsP3 membrane dynamics. *(A)* three-dimensional view (*ϕ* = 360*°, θ* = 20*°*) and *(B)* two-dimensional Mollweide projection. Each frame corresponds to 10sec in realtime. The mesh size is *m*_*max*_ = 1544. The cell volume and adhesion radius is 0.95*p*L and 4.5*μm*, respectively. The line indicates the interface between adhesion and non-adhesion membrane areas.

**Movie S2.**
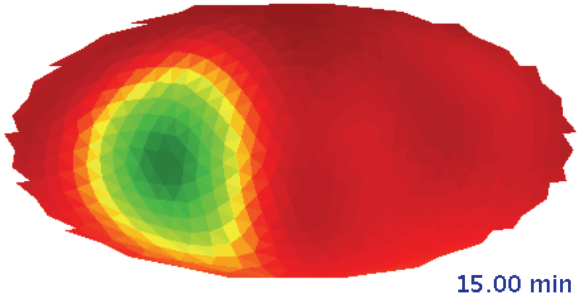
Animation of PtdInsP3 propagating wave dynamics along the equatorial periphery mapped on two-dimensional Mollweide projection. Each frame corresponds to 10sec in realtime. The mesh size is *m*_*max*_ = 1892. The cell volume and adhesion radius is 1.25*p*L and 2.6*μm*, respectively. The line indicates the interface between adhesion and non-adhesion membrane areas.

**Movie S3.**
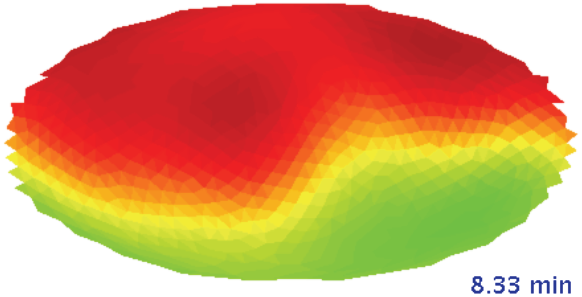
Animation of PtdInsP3 propagating wave dynamics along the latitude of the membrane mapped on two-dimensional Mollweide projection. Each frame corresponds to 10sec in realtime. The mesh size is *m*_*max*_ = 1388. The cell volume and adhesion radius is 0.89*p*L and 5.6*μm*, respectively. The line indicates the interface between adhesion and non-adhesion membrane areas.

**Movie S4.**
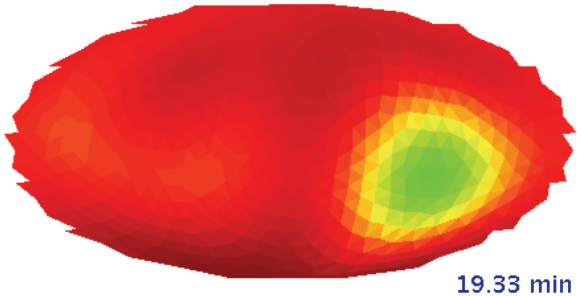
Animation of PtdInsP3 spot wave dynamics mapped on two-dimensional Mollweide projection. Each frame corresponds to 10sec in realtime. The mesh size is *m*_*max*_ = 956. The cell volume and adhesion radius is 0.53*p*L and 4.4*μm*, respectively. The line indicates the interface between adhesion and non-adhesion membrane areas.

**Movie S5.**
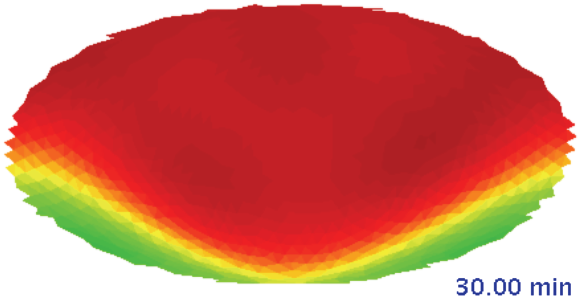
Animation of PtdInsP3 standing wave dynamics mapped on two-dimensional Mollweide projection. Each frame corresponds to 10sec in realtime. The mesh size is *m*_*max*_ = 1892. The cell volume and adhesion radius is 1.30*p*L and 5.1*μm*, respectively.

Author contributions
M.H. and T.S. designed research; M.H. performed data acquisition; M.H. implemented the analysis routine; M.H. and T.S. analyzed data; and M.H. and T.S. wrote the paper.

